# Analysis of the upper respiratory tract microbiota in mild and severe COVID-19 patients

**DOI:** 10.1101/2021.09.20.461025

**Authors:** V. Babenko, R. Bakhtyev, V. Baklaushev, L. Balykova, P. Bashkirov, J. Bespyatykh, A. Blagonravova, D. Boldyreva, D. Fedorov, I. Gafurov, R. Gaifullina, J. Galeeva, E. Galova, A. Gospodaryk, E. Ilina, K. Ivanov, D. Kharlampieva, P. Khromova, K. Klimina, K. Kolontarev, N. Kolyshkina, A. Koritsky, V. Kuropatkin, V. Lazarev, A. Manolov, V. Manuvera, D. Matyushkina, M. Morozov, E. Moskaleva, V. Musarova, O. Ogarkov, E. Orlova, A. Pavlenko, A. Petrova, N. Pozhenko, D. Pushkar, A. Rumyantsev, S. Rumyantsev, V. Rumyantsev, L. Rychkova, A. Samoilov, I. Shirokova, V. Sinkov, S. Solovieva, E. Starikova, P. Tikhonova, G. Trifonova, A. Troitsky, A. Tulichev, Y. Udalov, A. Varizhuk, A. Vasiliev, R. Vereshchagin, V. Veselovsky, A. Volnukhin, G. Yusubalieva, V. Govorun

## Abstract

The microbiota of the respiratory tract remains a relatively poorly studied subject. At the same time, like the intestinal microbiota, it is involved in modulating the immune response to infectious agents in the host organism. A causal relationship between the composition of the respiratory microbiota and the likelihood of development and the severity of COVID-19 may be hypothesized. We analyze biomaterial from nasopharyngeal smears from 336 patients with a confirmed diagnosis of COVID-19, selected during the first and second waves of the epidemic in Russia. Sequences from a similar study conducted in Spain were also included in the analysis. We investigated associations between disease severity and microbiota at the level of microbial community (community types) and individual microbes (differentially represented species). To search for associations, we performed multivariate analysis, taking into account comorbidities, type of community and lineage of the virus. We found that two out of six community types are associated with a more severe course of the disease, and one of the community types is characterized by high stability (very similar microbiota profiles in different patients) and low level of lung damage. Differential abundance analysis with respect to comorbidities and community type suggested association of *Rothia* and *Streptococcus* genera representatives with more severe lung damage, and *Leptotrichia*, unclassified Lachnospiraceae and *Prevotella* with milder forms of the disease.

## 1 Introduction

Commensal microorganisms of the respiratory tract affect the body’s ability to withstand the threat of viral infections [1] [2]. This effect is due to the modulatory influence of microbes on the immune system [3] or direct interaction between microbial metabolites with the viral life cycle [4].

At the same time, certain bacteria can promote viral infections [5] and bacterial secondary infections are leading to serious complications and mortalities during viral infections including new coronavirus disease [6][7][8]. Viral infections in turn can influence the bacterial community, sometimes promoting bacterial biofilm formation and secondary infections [9][10]. The close relationship between respiratory microbiota and viral infections renders it an important subject of study [11].

Until recently the microbiota of the upper respiratory tract was poorly studied, especially in adults [12]. This situation is changing in the context of the current coronavirus pandemic. Previous studies identified the existence and dynamical nature of microbial community types of upper respiratory tract during COVID-19 infection [13]. A number of studies described the differences between patients with COVID-19 and healthy people or differences among patients with different disease severity ([14][15] among others), see [16] for a short review. The results of different studies are quite different to each other which may be the result of phenotypic differences among groups studied (e.g. comorbidities), season and geographical factors and different techniques used in the data sampling and analysis.

Different approaches to study microbiota associations are used by different authors when analysing current COVID-19 pandemic samples. In [13] authors analyze throat and fecal specimens from 35 COVID-19 patients, 19 healthy adults and 10 non-COVID-19 patients. They obtained amplicon sequence variants (ASV) using DADA2 method, utilized Dirichlet multinomial mixtures probabilistic modelling and infer six community types, found associations between infection status and community type and also examined the change of type in the course of coronavirus disease. The majority of COVID-19 patients had microbial respiratory community type 2, characterized by higher abundance of *Porphyromonas*, *Neisseria*, *Fusobacterium* and Bacteriodales. The synchronous change of community types in the upper respiratory tract and gut was observed in a number of patients. In [14] authors examined 38 COVID-19 patients and 21 uninfected controls. They obtained amplicon sequence variants using DADA2 method, and utilized DESeq2 to infer ASVs associated with the COVID-19 and also with higher viral load. It was observed that *Peptoniphilus lacrimalis*, *Campylobacter hominis*, *Prevotella copri*, and an *Anaerococcus* unclassified amplicon sequence variant were associated with the disease and higher viral load, whereas *Corynebacterium* unclassified, textitStaphylococcus haemolyticus, *Prevotella disiens*, and two *Corynebacterium 1* unclassified amplicon sequence variants were associated with healthy status and low viral load during COVID-19.

Respiratory microbiome has geographic and climatic characteristics, so it is important to study its relationship with COVID-19 in many regions. It is also essential to take into account other diseases and habits of patients when analyzing the associations between the composition of the microbiota and the severity of the course of covid disease, to lower the rate of spurious correlations.

In our work, we describe the composition of the respiratory microbiome in 219 inpatients and 146 outpatients with COVID-19. We performed SARS-CoV-2 genome sequencing and metagenomic analysis of nasopharyngeal swabs to find associations with the disease severity. As far as we know, the current paper is the first description of the respiratory tract of patients with COVID from Russia and one of the few studies in which microbiome analysis was performed regarding patient’s comorbidities.

Two approaches were used in metagenomic data analysis. First we inferred groups of microbial communities using Dirichlet multinomial mixtures (DMM) probabilistic modelling. Groups of microbial communities unite communities with similar taxonomic profiles, and modeling of taxonomic profiles as the result of multinomial sampling addresses the compositional nature of metagenomic sequencing data [17]. This method gained popularity in microbiome research in recent years [18][19][13].

Secondly we aimed at identification of microorganisms associated with more severe forms of disease using differential abundance testing with the use of patient’s metainformation as covariates. Microbial community type was also used as covariate to separate the effects on the community level as a whole and the influence of individual microorganisms. We performed testing with the DESeq2 [20] method which demonstrates sensitivity and low false positive rate in the microbiome data analysis [21].

We found that two out of six community types are associated with a more severe course of the disease, and one of the community types is characterized by high microbial content stability and low level of lung damage in patients. Differential abundance analysis suggested association of *Rothia* and *Streptococcus* genera representatives with more severe lung damage, and *Leptotrichia*, unclassified Lachnospiraceae and *Prevotella* with milder forms of the disease. The wide geographical and age coverage of COVID-19 patients with taking into account their comorbidities allows us to hope for further reproducibility of the obtained results.

## 2 Results

### 2.1 Study cohort

336 patients were enrolled in the study aged up to 94 years, 203 females and 131 males, 140 outpatients and 196 inpatients (the total number of patients and the number of patients in the groups will not match due to some missing data). Samples were collected between 2020-05 and 2021-03 (collection dates capture the decay of wave 1 and the entire period of wave 2 of covid 19 incidence) in the following cities of Russian Federation: Moscow, Kazan, Irkutsk, Nizhny Novgorod (Figure 1).

**Fig. 1.**
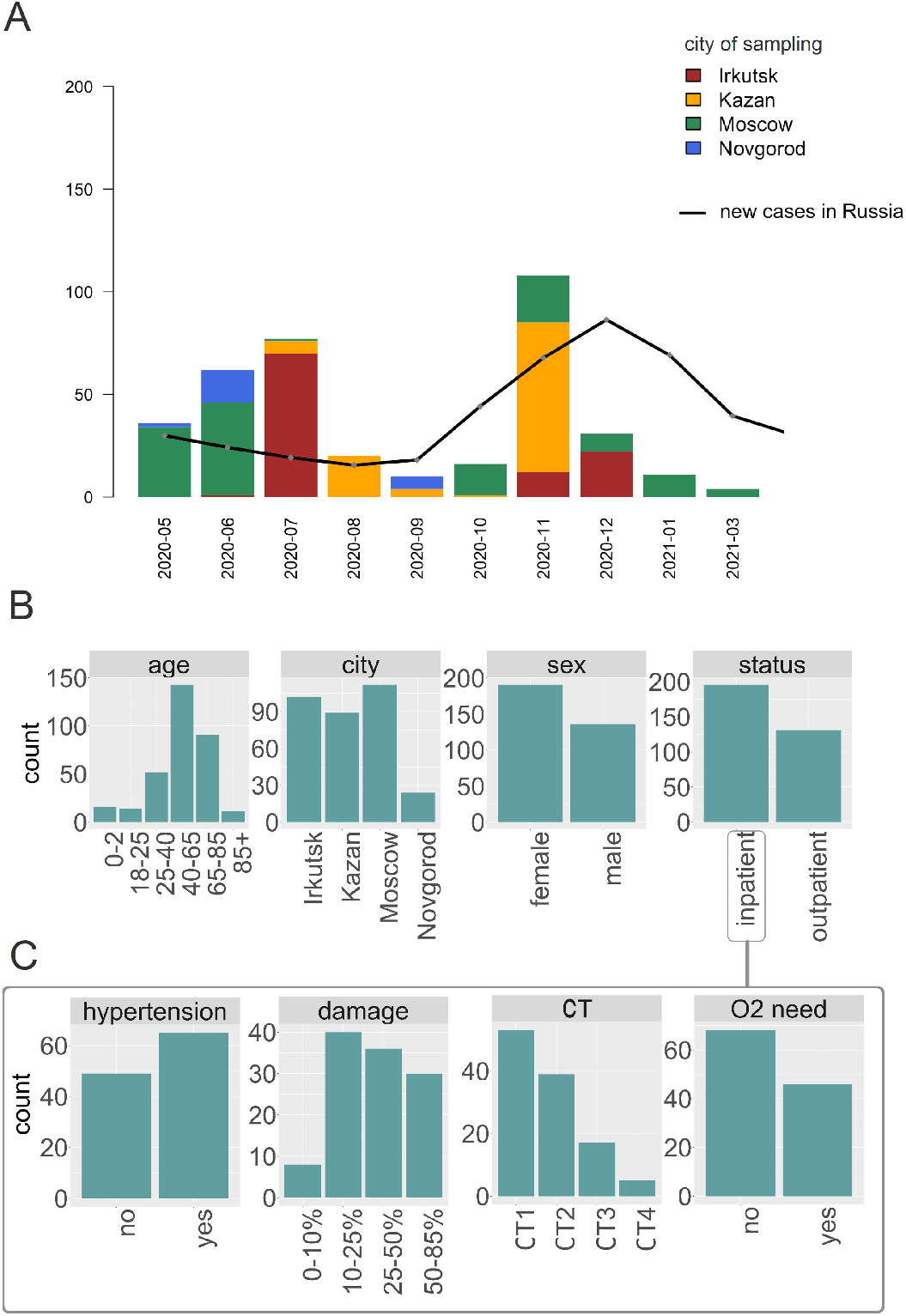
Study cohort overview. A) The number of samples collected in different months and in different cities. The black line shows the number of newly identified SARS-CoV-2 infection cases in Russia. B) distribution of patients by age, sex, city of sampling, hospitalization status for all patients included in the study. C) The clinical characteristics of inpatients are shown: the percentage of lung damage, computer tomography severity score (CT), the need for additional oxygen supply as well as the presence of hypertension as a concomitant disease.

Data on both inpatients and outpatients were obtained for the city of Irkutsk. For the city of Kazan, we had only outpatients and data on them was used only when analyzing community types. The following clinical information was collected: computer tomography severity score (CT), percent of lung tissue affected, temperature, oxygen saturation level, respiratory rate, heart rate, systolic and diastolic blood pressure, being conscious and the need for additional oxygen supply. The following metadata was collected based on questionnaires: presence of obesity, diabetes, chronic obstructive pulmonary disease (COPD), inflammatory bowel disease (IBD), arthritis, tuberculosis, hypertension, coronary artery disease, chronic heart failure, asthma, current and past smoking status.

### 2.2 Analysis of comorbidities and habits associated with the disease severity

We analyzed the association of various available clinical and non-clinical parameters with the proportion of affected lung tissue. The presence of hypertension was significantly associated with linear model analysis (p-value < 0.001) which is in agreement with previous reports [22]. No other associations of disease severity and patient data and comorbidities were found, presumably due to the relatively small number of patients. No associations of disease severity and virus genome lineage (pangolin classification) were found to be statistically significant.

### 2.3 SARS-CoV-2 genome sequences

We sequenced genomes of SARS-CoV-2 virus amplified from the patient’s swabs. The coverage width of the 258 genome sequences was higher than 90

Most of the genomes belonged to B.* pangolin lineages, namely: B.1.1 (183 sequences, found all over the world), B.1.1.317 (26 sequences, prevalent in Russia and other European countries), B.1 (16 sequences, found all over the world), B.1.1.294 (12 sequences, prevalent in Russia and other European countries), B.1.1.163 (7 sequences, sampled mainly in Russia); one genome belonged to M3 lineage, which is mainly sampled in England.

We observed significant association between lineages and the city of sample collection (Fisher test with simulated p-values, p-value < 3 × 10^−5^). Uneven prevalence was observed mostly for lineage B.1.1.294 which was quite frequently observed in Kazan (exact Fisher test, p-value < 0.002).

No association was found between pangolin lineage of the virus genome and CT score or percentage of lung damage (linear models analys).

### 2.4 Respiratory microbiome analysis

The 16S-rRNA gene sequences were resolved into 3341 amplicon sequence variants (ASVs) representing 345 genera. Further we will describe the analysis of community types, analysis of factors associated with community types and the association of community types with clinical indicators and will conclude with differential abundance analysis on the genus and ASV levels.

#### 2.4.1 Microbial community types analysis

Microbial community types analysis We observed the presence of several clusters of taxonomic profiles in our data (see Figure 2). We analyzed jointly our data and data from work [23] to infer community types using Dirichlet multinomial mixtures (DMM) modelling. A joint analysis was performed to increase confidence in the resulting clusters. We got 6 community types based on the Laplace approximation criteria (Figure 2 A,C).

**Fig. 2.**
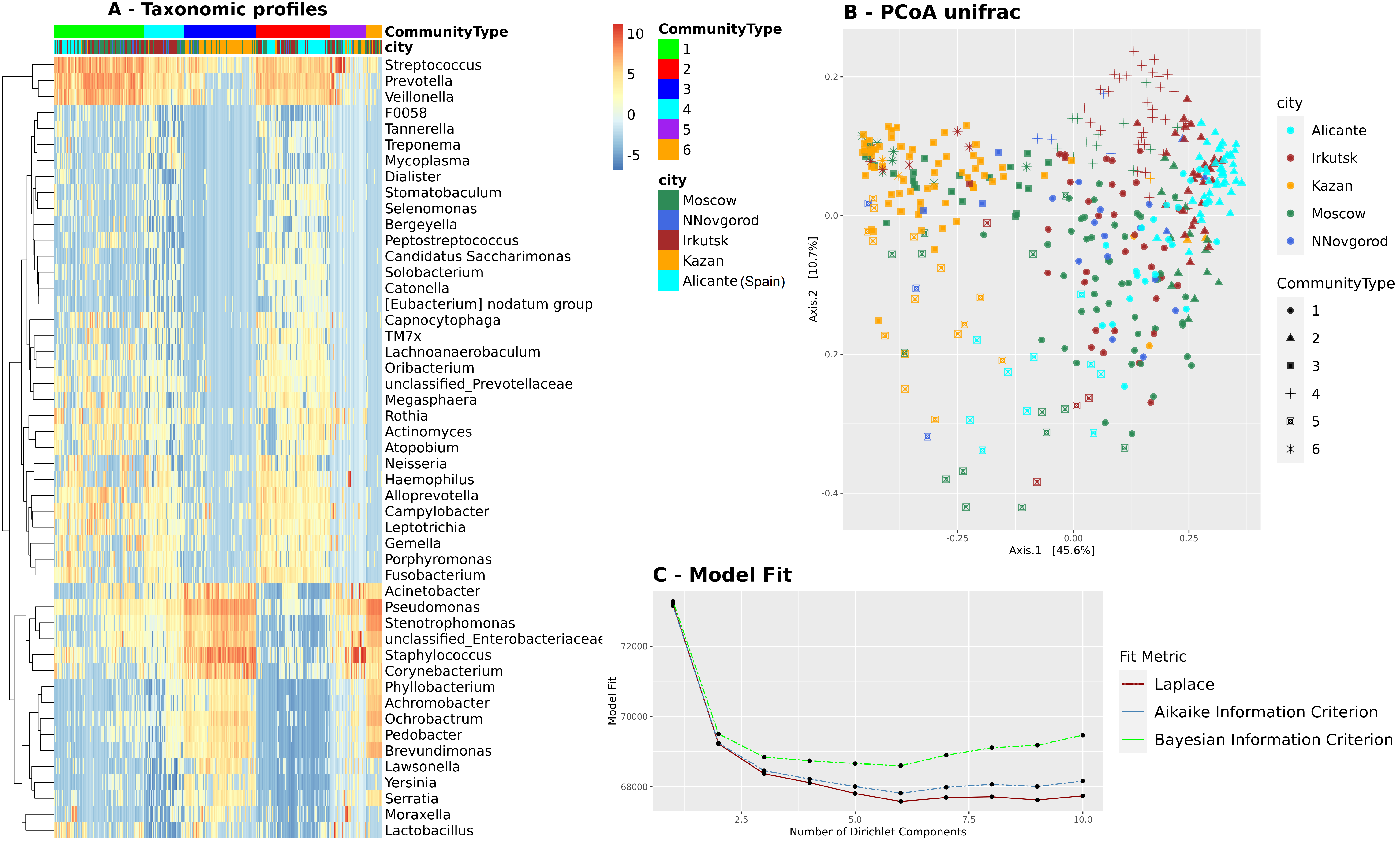
Microbial community types analysis. A) Heatmap visualization of taxonomic profiles on the genus level with inferred community type and city of sampling designated with color; Alicante city corresponds to external samples obtained in the study [10.3389/fmicb.2021.637430]. B) PCoA diagram based on unifrac distances between samples; color shows the city from which the sample was collected, the form - the type of community. C) Cluster analysis using DMM modeling infers six types of microbial communities based on the Laplace approximation.

Supplementary Figure 2 shows estimated relative abundances of different microbial genus in six community types.

According to our analysis, the main drivers of community types 1 and 4 are *Streptococcus*, *Prevotella* and *Veillonella* genera, but community type 4 is characterized by larger alpha diversity (Figure 4 C). Community type 2 is similar to them and has the same top three drivers, but *Prevotella* prevails over streptococci. Community types 3 and 5 are also similar to each other with the most represented genera being *Staphylococcus*, *Pseudomonas* and *Corynebacterium* for type 3 and *Staphylococcus*, *Pseudomonas*, *Streptococcus* for type 5. Community type 6 is mainly dominated by *Pseudomonas* and *Stentotraphomonas* genera. The communities are numbered in descending order of the number of samples assigned to them.

**Fig. 3.**
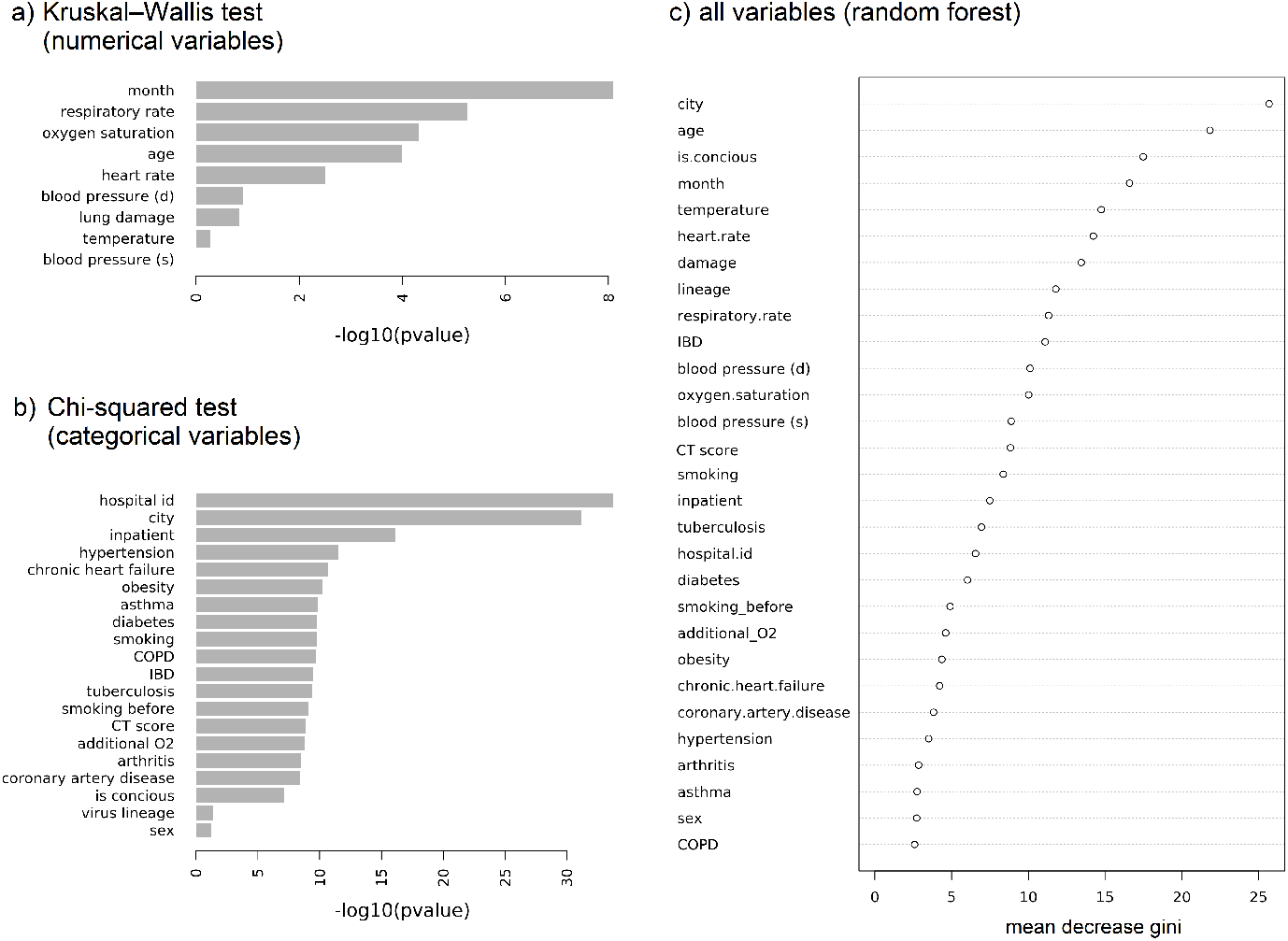
Factors associated with the type of microbial community of the upper respiratory tract. The negative logarithm of p-value is shown for the Kruskal-Wallis test for numerical variables (a) and in the chi-square test for categorical variables (b); the association between the individual variable and the type of community was estimated, the larger the value, the more significant the association. C) the importance of variables (mean decrease Gini index) in the random forest is shown. Month, city and hospital id concerns the sampling of the material; age, temperature, oxygen saturation, respiratory rate, heart rate, blood pressure (diastolic) and blood pressure (systolic), being conscious (is concious) and being hospitalized (inpatient), the need for additional oxygen supply (additional O2) are the factors describing the patient’s condition; obesity, smoking, smoking before, diabetes, COPD (chronic obstructive pulmonary disease), IBD (inflammatory bowel disease), arthritis, tuberculosis, hypertension, coronary artery disease, chronic heart failure, asthma are concomitant diseases of the patient and his habits, lineage is the SARS-CoV-2 genome Pangolin classification.

**Fig. 4.**
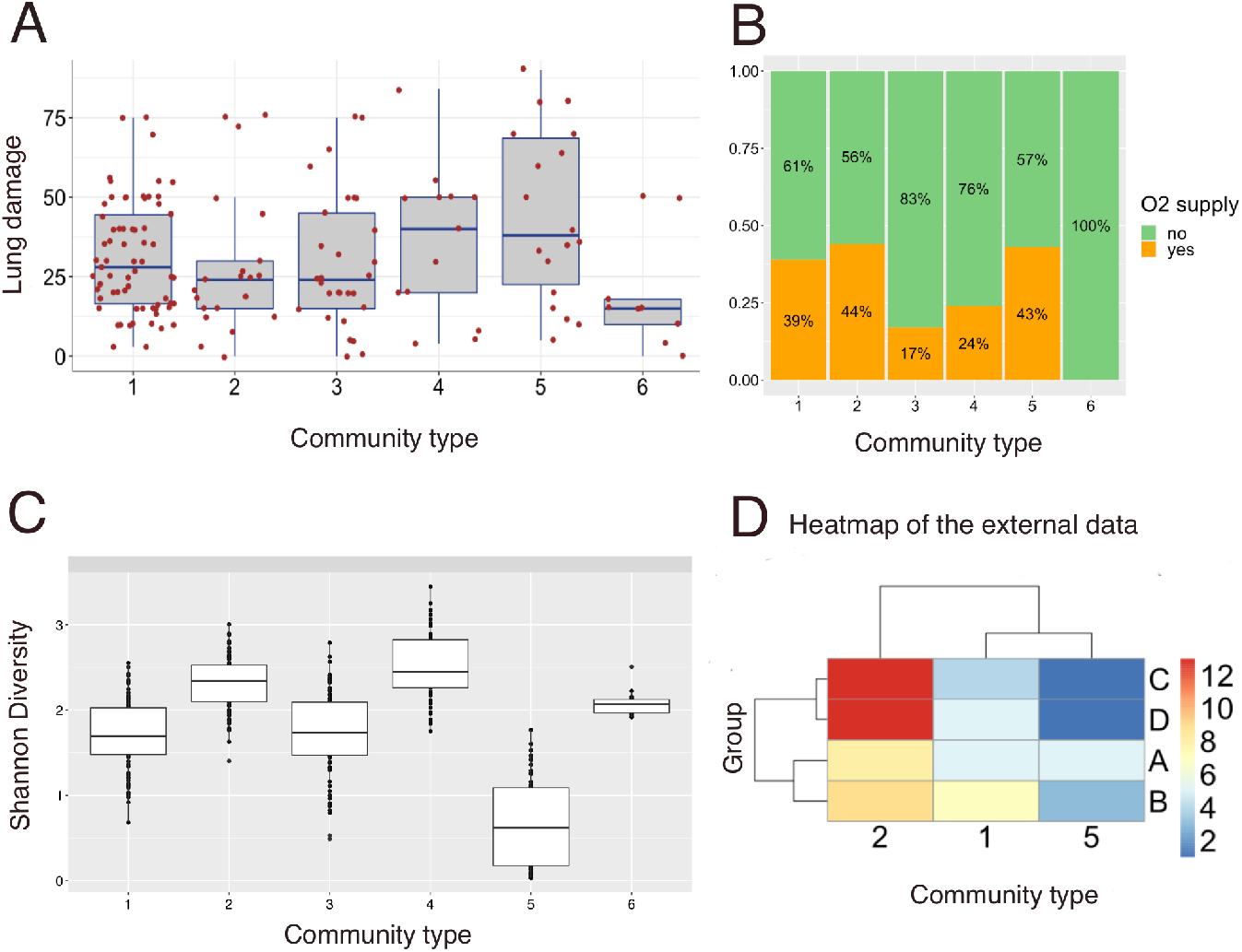
The association between the microbial community type and the severity of the disease. The distribution of the proportion of lungs affected (A) and the need for additional oxygen supply (B) by community types is shown. The alpha diversity (measured as Shannon index) of different community types (C). Section D shows the distribution by type of communities of patients from the study [23]; groups show the severity of the disease (group A: SARS-CoV-2 negative patients (n = 18); group B: mild COVID-19 symptoms but no later hospital admission (n = 19); group C: severe COVID-19 symptoms followed by hospital admission (n = 18); and group D: patients with severe COVID-19 symptoms which were eventually admitted into intensive care units (n = 19))

**Fig. 5.**
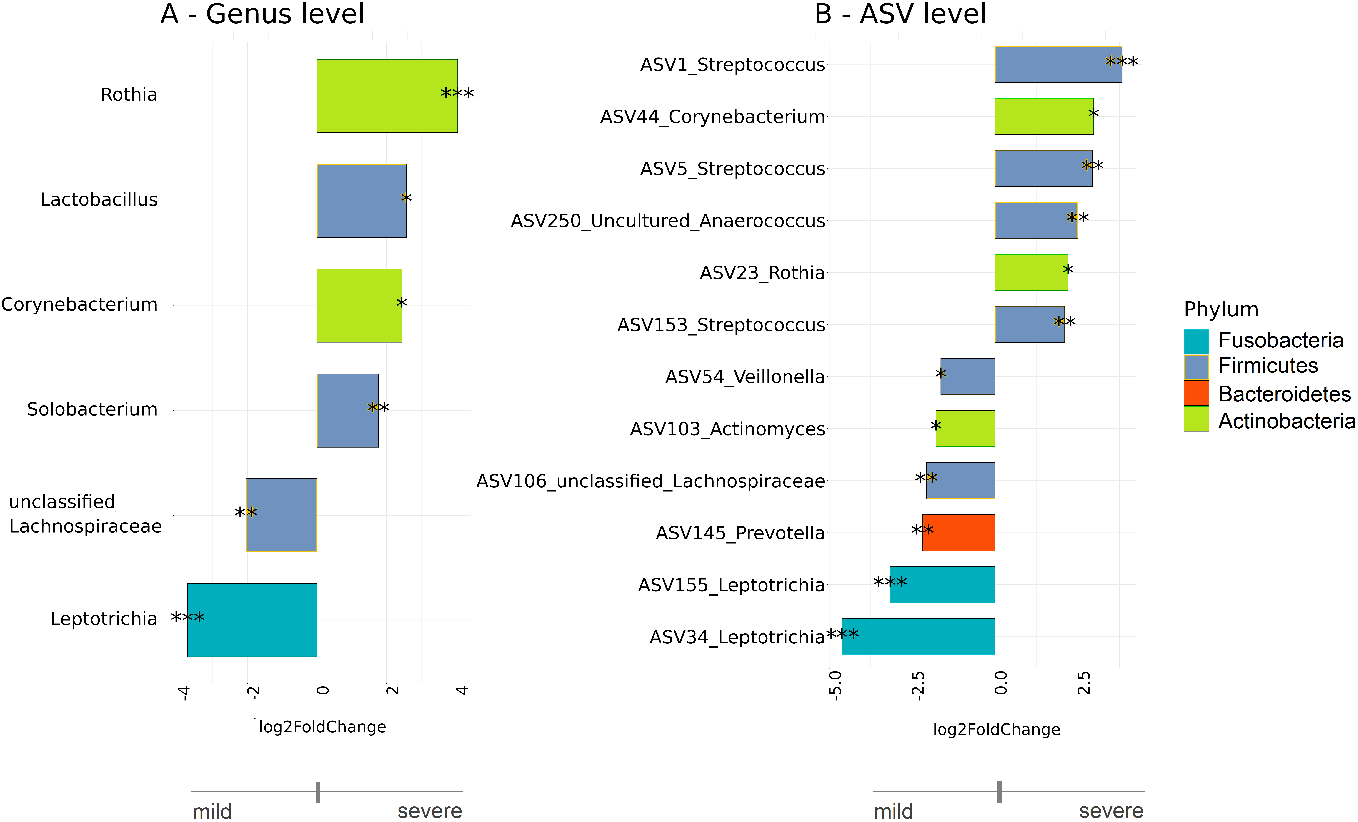
The differentially abundant taxa testing associations. Bacteria on the genus (A) and ASV (B) levels that differ in abundance in inpatients with severe (CT ¿ 2) and mild (CT = 1) level of lung damage. The analysis was carried out using the DeSeq2 method. Bacteria which are more abundant in more severe patients are shown in bars pointing to the right, in lighter patients - to the left. Asterisks indicate the level of significance with the following thresholds: (*) = 0.1, (**) = 0.05, (***) = 0.01.

#### 2.4.2 Factors affecting the type of community

We analyzed the association between the variables characterizing the patient and his or her condition and the microbial community type using the following approaches. First, we ran the Kruskal-Wallis nonparametric statistical test for each of the numerical variables (Figure 3 A) and the chi-squared test for each of the categorical variables (Figure 3 B). We also applied the random forest method to estimate the importance of both categorical and numerical variables (Figure 3 C).

Different methods give different results but their consensus is that city and month of sample collection and patient’s age are the most significant variables associated with the microbial community type. The association between age and the composition of the upper respiratory tract microbial community is rather well known [24]. The influence of season of sample collection on microbiota composition was previously shown for children [25]. Air microbiome was shown to be season and geographical location dependent in [26] with Pseudomonas genera being one of the most variable community members (see Figure 3 from [26]).

As shown by post hoc analysis, community types 2 and 4 are associated with an earlier age of the patients (Supplementary Figure 1 A, statistical analysis performed with dune’s test) with community types 2 and 4 containing younger people than other types. The strongest association with the month of sample collection is observed in community type 3 (Supplementary Figure 1 B), samples belonging to this type were collected mainly in winter.

Different cities have different distributions of community types (Fisher exact test with simulated p-values for all possible pairs of cities, FDR adjusted p-value < 0.05), with the only exception being the comparison of Moscow and Novgorod, the difference between which is not significant. As shown in Supplementary Figure 1C in Moscow and Nizhniy Novgorod predominates community type 1; in Kazan city type 3 is more prevalent and in Irkutsk city a relatively uniform distribution of types 1, 2 and 4 is observed. Samples from [23] which were collected in Alicante belonged mainly to community types 2 and 1.

#### 2.4.3 Association of microbial community types with disease severity

We observed a statistically significant difference in the need of additional oxygen supply for patients with microbiota of different types (p-value < 0.003, Fisher exact test). There were some differences in the degree of lung damage, but it was not statistically significant (Kruskall-Wallis test, p-values ¿ 0.05). As a trend, we observe a larger lung damage and larger need in O2 supply in community type 5, which also shows the lowest alpha-diversity (Figure 4 A, B, C).

As well, the high proportion of patients with additional oxygen needs has type 2 microbial community. Interesting that the majority of patients with a severe form of the COVID-19 from the [23] study also belonged to the second type of community.

A comparison of in- and outpatients from the city of Irkutsk (the only city where we managed to collect samples of both types) showed that the microbiota of inpatients belongs mainly to community types 1 and 2, and the microbiota of outpatients belongs mainly to type 4. The type 4 community had the highest level of alpha diversity and patients with this type of community rarely required additional oxygen supply while having sufficiently high levels of lung damage.

As was mentioned above, in our study the 5th type of community having the extremely low level of alpha diversity was predominant in patients with a high percentage of lung damage and often needed oxygen supply. Otherwise in [23] this type was found mainly in patients belonged to group A: SARS-CoV-2 negative patients (n = 18) and group B: mild COVID-19 symptoms but no later hospital admission (n = 19) (Figure 4D).

The 6th type of community was the most stable in its composition (figure 2 A). Patients having this type of community were sampled in different cities, had no need for additional oxygen supply, and had a low level of lung damage.

#### 2.4.4 Microbial associations of disease severity with respect to community type, season and comorbidities

To search for the potential effect of microbiome composition on the severity of COVID-19, we took into account the following metadata: age, sex of the patient, presence of hypertension, city and season of the sampling, microbial community type and the SARS-CoV-2 virus lineage. We analyzed the differentially abundant genus, between patients with mild (CT 1) and more severe (CT 2 - 4) levels of lung damage, using the DESeq2 method [20].

It may be noted that in patients with mild lung damage (CT 1), significantly (p-value < 0.01) predominates bacteria belonging to Fusobacteria phylum which are known as commensals of the upper respiratory and gastrointestinal tracts in humans and animals. In particular, based on analysis on the level of amplicon sequence variants (ASV) we observed the overrepresentation of bacteria from the *Leptotrichia* genus, especially ASV34 and ASV155, which have identical sequences with *Leptotrichia wadei* and a number of uncultured bacteria, respectively.

*Leptotrichia* species are anaerobic or facultative anaerobic species frequently found in the oral cavity and are able to produce lactic and some other organic acids [27]. We are unaware of the possible positive biological role of *L. wadei* and uncultured *Leptotrichia* species in promoting milder disease. It can also be assumed that the upper respiratory tract microbiota characteristic for healthy people remain in the group of patients with mild lung damage.

The *Prevotella* genera and unclassified Lachnospiraceae (both on the genus and ASV levels) also appeared to be associated with mild lung damage. Along with *Leptotrichia*, *Prevotella* species belonging to Bacteroidetes phylum are mainly known as commensals in the respiratory tract and rarely cause respiratory infections [28]. Depletion of *Prevotella* was observed in patients with influenza infection [29][30] and also in a group of patients with different acute viral infections (influenza A, influenza B, rhinovirus, metapneumovirus and respiratory syncytial virus) [31]. But *Prevotella* spp. produced proteins were found to be abundant in the lung metagenomic studies of SARS-CoV-2 infected patients [32].

The Lachnospiraceae family is a phylogenetically and morphologically heterogeneous taxon belonging to the Firmicutes phylum. It is considered to be a member of the core gut microbiota, but also found in bronchoalveolar lavage and sputum. Bacteria from the Lachnospiraceae family are able to ferment sugars and produce butyrate and other short-chain fatty acids (SCFAs) in the gut [33]. SCFAs are known to facilitate gut barrier function and to modulate the immune system with mostly beneficial effects [34], including protection against respiratory viruses [35]. These substances also play a role in a gut-brain axis [36]. The role of SCFAs in the respiratory tract is less studied, while they can be found in bronchoalveolar lavages [37] and potentially can affect airway inflammatory responses [38][39]. Previously, it was shown that the presence of the Lachnospiraceae in bronchoalveolar lavage samples predicted worse ICU outcomes for critically ill patients [40]. According to our data it seems that Lachnospiraceae somehow protect the people with COVID-19 from developing the severe lung damage, but it should be analyzed in more detail in the future studies.

Among the patients with severe lung damage (CT2-4) we observed the overrepresentation of bacteria from *Rothia* (p-value < 0.01) and *Solobacterium* (p-value < 0.05) genera. *Rothia* species (*R. mucilaginosa*, *R. dentocariosa*, *R. aeria*, *R. nasimurium*, and *R. amarae*) being a part of the normal flora of the human oropharynx and upper respiratory tract, can cause a wide range of serious infections, including pneumoniae, especially in immunocompromised people. Earlier bacteria from the *Rothia* genus have been already observed in association with COVID-19. In [13] this genus was described as a major taxon in one of the throat microbial community types represented in patients but not in healthy controls. In another study, ASV identical to 16S rRNA of *Rothia dentocariosa* was found to be overrepresented in samples from COVID-19 patients [41]. There is also at least one published work describing the lower abundance of *Rothia* species in COVID-19 patients [42]. The analysis performed on the ASV level showed that three ASVs from the *Streptococcus* genus and one ASV of the *Anaerococcus* genus are overrepresented in patients with more severe lung damage.

Three Streptococci ASVs associated with greater lung damage have exact sequence match with *Streptococcus mitis/oralis* (ASV1), *Streptococcus parasanguinis* (ASV5) and *Streptococcus anginosus* (ASV153) 16S rRNA sequences. These species are representatives of viridans group streptococci having mainly commensal relationships with the human host, but also capable of causing or promoting respiratory infections [43]. *Oral/mitis* streptococci were previously described as potentially contributing to influenza viral infection [4], supernatants of these bacteria promoted viral release and spread presumably due to neuraminidase production by these bacteria. *Streptococcus anginosus* plays an important role in respiratory infections [44], including bacterial pneumonia in COVID-19 patients [45]. The abundance of *Streptococcus* genera and particularly of *S. parasanguinis* on hospital admission, was identified as a strong predictor of fatality in COVID-19 patients [**?**].

## 3 Summary

Our study aimed at characterization of nasopharyngeal microbiota characterization of COVID-19 patients with different levels of disease severity.

First, we performed analysis on the level of community types. Mixture model analysis resulted in six types of bacterial communities. Among the factors associated with community type are geographic location and month of sample collection and the age of the patient. As a trend, patients with community type dominated by *Pseudomonas* and *Stentotraphomonas* genera had relatively low lung damage and no need for additional O2 supply. Patients with low alpha-diversity community type with *Staphylococcus*, *Pseudomonas*, *Streptococcus* as top genera demonstrated the highest lung damage.

Next we performed the analysis of differentially abundant taxonomic units with respect to confounding parameters, including respiratory community type, inferred *Rothia* and *Solobacterium* genera and sequence variant of *Streptococcus mitis/oralis* group to be positively correlated with larger lung damage levels. *Leptotrichia* and unclassified Lachnospiraceae genera, sequence variants belonging to *Prevotella* genus were observed to be associated with lower lung damage.

## Supporting information

Supplementary figure 2. Estimated relative abundance of different microbial genus in six community types.

Supplementary figure 1. Distribution of the patient's age and the month of sampling between different community types.

## Ethics approval and consent to participate

The study was approved by the ethical committee of RCPCM. All patients gave written informed consent for sample collection and personal data processing.

## Acknowledgments

We thank the Center for Precision Genome Editing and Genetic Technologies for Biomedicine, Federal Research and Clinical Center of Physical-Chemical Medicine of Federal Medical Biological Agency for providing computational resources for this project.

