## Supplementary figure 2. Estimated relative abundance of different microbial genus in six community types. for "Analysis of the upper respiratory tract microbiota in mild and severe COVID-19 patients"

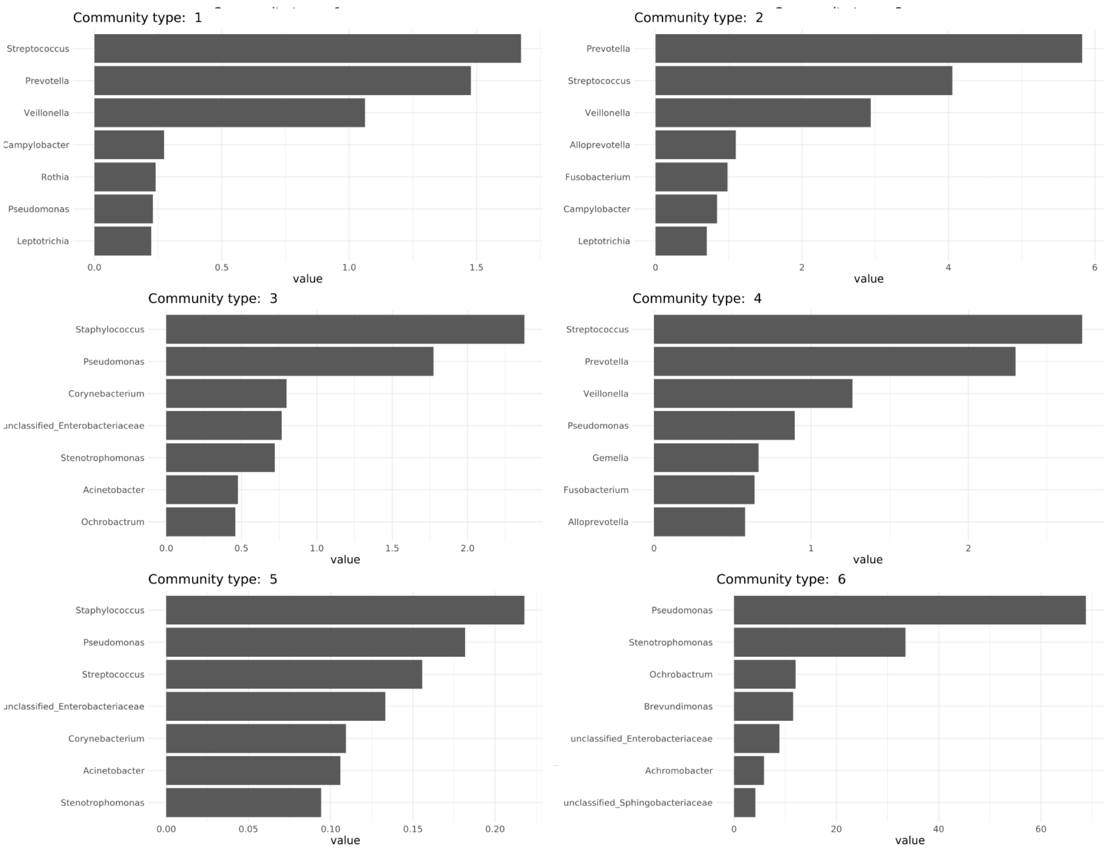


Supplementary figure 2. Estimated relative abundance of different microbial genus in six community types.
