## Supplementary figures and images for "Analysis of the upper respiratory tract microbiota in mild and severe COVID-19 patients"

### Supplementary figure 1. Distribution of the patient's age and the month of sampling between different community types.

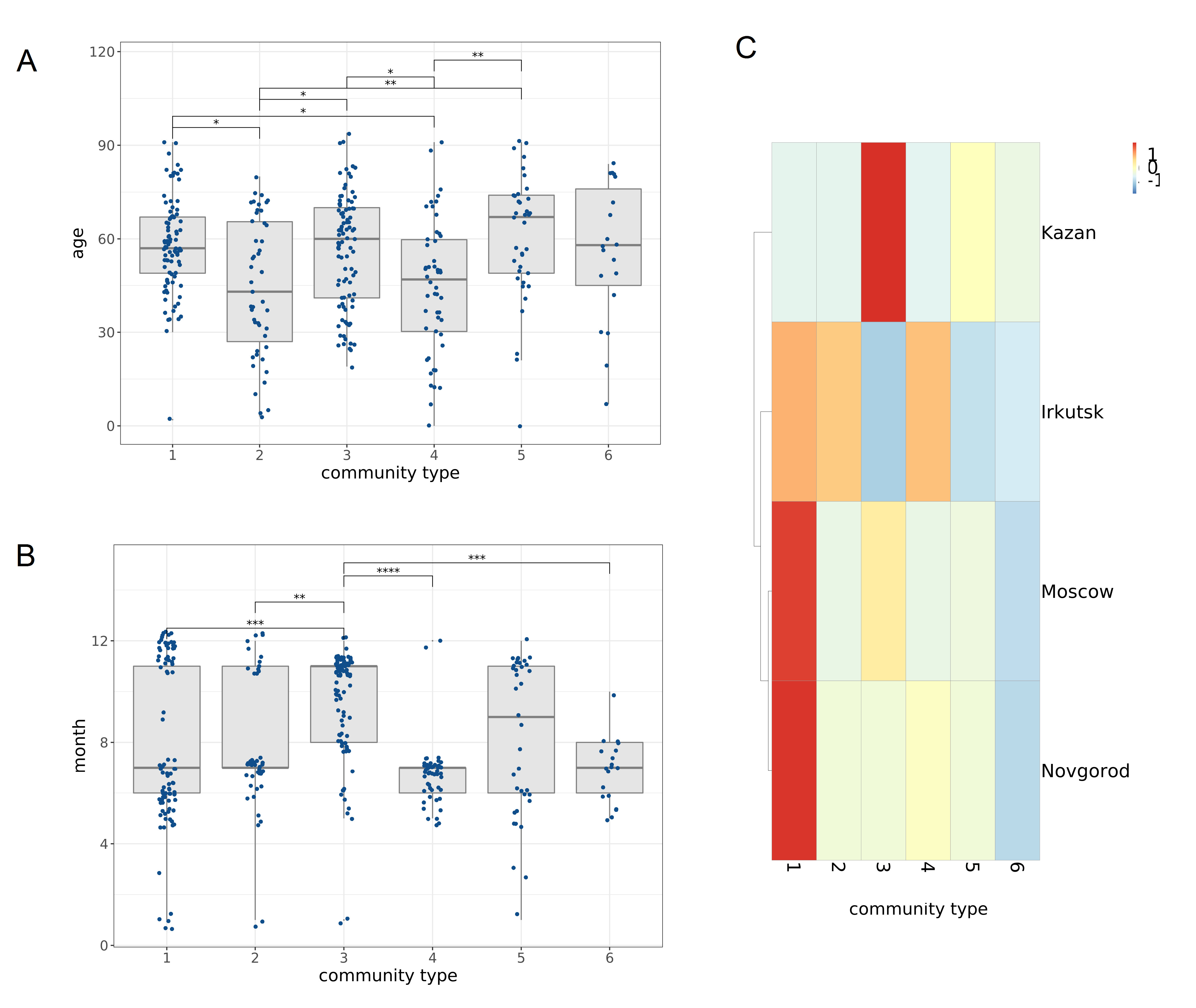
